# Embedding the Skin Conductance Response into the Brain Connectivity Framework: Monoaminergic Signaling Visible Through the Lenses of Computational Modeling

**DOI:** 10.1101/504183

**Authors:** Saša Branković

## Abstract

Relying on evidence for the functional, neurochemical, and spectral parallelism between the late event-related potentials, delta oscillatory brain responses, and the skin conductance response (SCR) system the hypotheses about the existence of the SCR-related brain oscillations and their connectivity with the SCR system have been here suggested. In contrast to classical approach to event-related oscillations which relies on either stimulus- or response-locked time reference, an approach assigned as “oscillatory process-related oscillations” has been introduced. The method enables us to overcome the variability of latency period of the SCR. The hypothesis about the existence of the SCR-related brain oscillations and their delta nature has been confirmed through the grand averaging method. An unexpected finding was the complex nature of the SCR-related oscillations: in addition to the two second EEG segment which was correlated with the SCR system signals they also comprised an initial 200 ms segment uncorrelated with the SCR. The hypothesis about the connectivity between the SCR system and the respective delta brain oscillatory response has been operationalized through a multiple time series regression model. The predictor set consists of the SCR, its first three derivatives, and their mutual interactions. The Monte Carlo test of the causal link between the SCR system signals and the related delta EEG signal demonstrated significance in more than half of the participants. The findings have been considered from the standpoints of the segmental structure of the EEG, monoaminergic signaling and recently emerged the “brain-body dynamic syncytium” hypothesis.

## Introduction

Theorizing on the neural basis of several, both central and peripheral psychophysiological measures in the last decade has implied the brainstem monoaminergic systems (dopaminergic, noradrenergic, and serotonergic). Dopaminergic neurotransmission has been for instance associated with the scalp-recorded P3a subcomponent of event related brain potentials (ERPs) and noradrenergic activity of the locus coeruleus (LC) with the P3b subcomponent (Polich 2007). According to Nieuwenhuis, Aston-Jones & Cohen (2005) P3 wave is the electrophysiological correlate of the LC-induced phasic excitation in the neocortex. Further, there is evidence (Labuschagne, Croft, Phan & Nathan 2010) that serotonin influences the late positive potential (LPP) in electrophysiological responding.

On the other hand, a peripheral autonomic signal, pupil dilation, has been found to indicate LC-noradrenergic activity (Hoeks & Ellenbroek 1993; Preuschoff, ’t Hart & Einhäuser 2011; Lavin, San Martin & Jubal 2014). Also, the skin conductance response (SCR) signal and its derivatives have been assocciated with the brainstem dopaminergic, noradrenergic, and serotonergic signaling (Branković 2008, 2012). Finally, the central EEG phenomena (ERPs) and peripheral autonomic components of orienting response (pupillary dilation and SCR) were supposed to be “intimately coupled” through the brainstem monoaminergic signaling (Nieuwenhuis, De Geus & Aston-Jones 2011; Branković 2014).

Since the brainstem monoaminergic signaling appears as a junction of these models of central and peripheral psychophysiological variables (Figure 1), a theoretical possibility emerges that there is connectivity between central and peripheral psychophysiological signals. Clinically even more interesting is a possibility that the supposed connectivity could reflect different facets of the monoaminergic functioning. Exploration of the existence and biological nature of the supposed connectivity could further contribute to the field of assessment of the brain neurochemistry through psychophysiological probing and computational modeling (Branković 2008, 2011, 2012).

**Figure 1.**
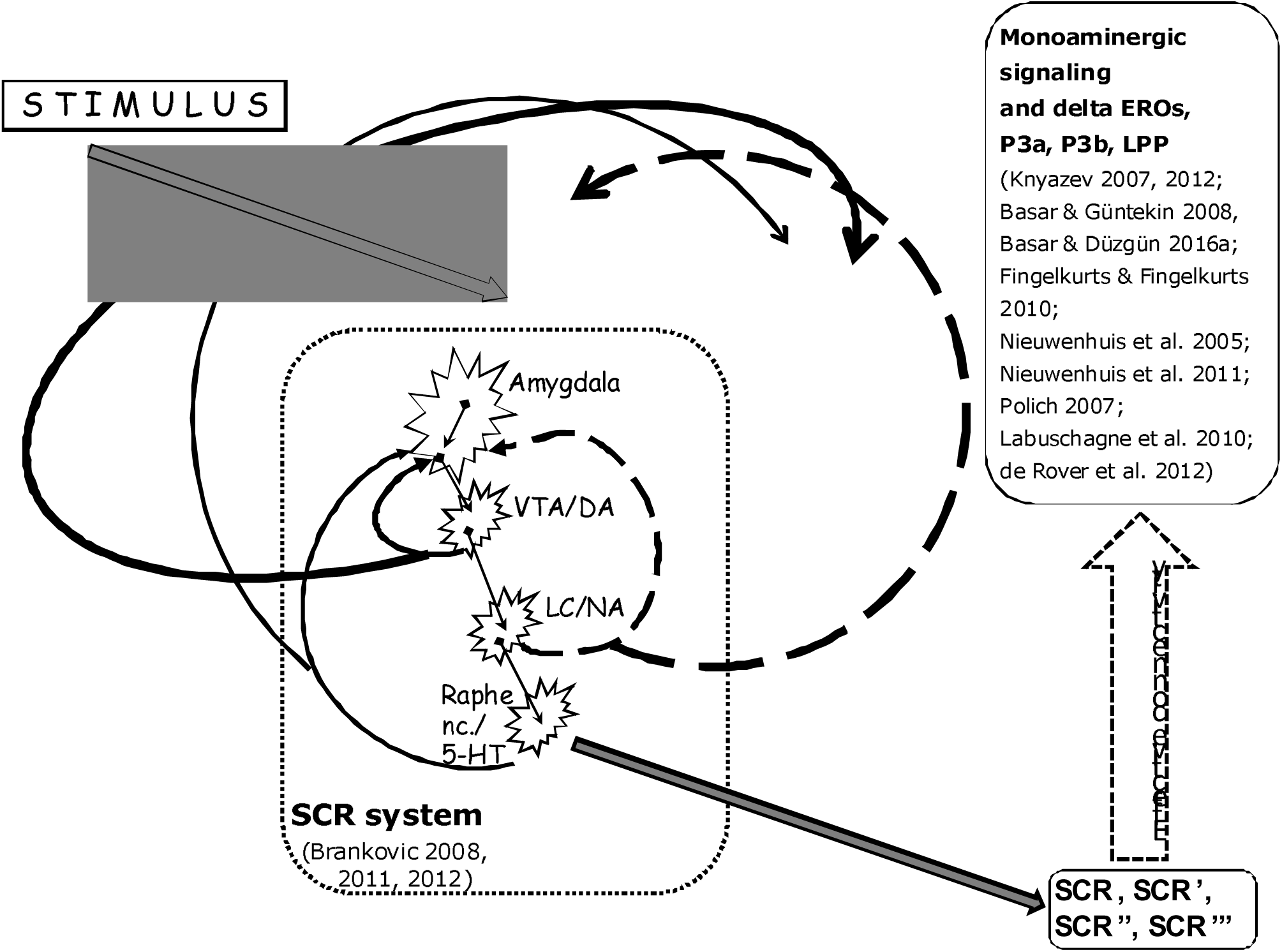
Monoaminergic signaling as a link between the late ERPs – delta oscillatory brain responses and the SCR system.

Here we deal with the hypotheses about (i) the existence of the SCR-related brain oscillations and (ii) the connectivity between the EEG signal and the SCR system. Before we expose the hypotheses in more details we review the two theoretical conceptualizations which are fundamental for the present approach. Those are (i) spectral and functional equivalence and feedback loops’ origin of the late event-related brain potentials and delta oscillatory brain responses (e.g. Knyazev 2007, 2012; Nieuwenhuis, Aston-Jones & Cohen 2005; Polich 2007; Başar & Güntekin 2008; Başar & Düzgün 2016a; Fingelkurts & Fingelkurts 2010) and (ii) the central nervous system interpretation of the mathematical model of the SCR (Branković, 2008, 2011, 2012).

### Spectral and functional equivalence and feedback origin of the later ERP components (P300 and LPP) and delta EEG responses

On spectral equivalence between P300 and delta responses has been pointed for more than thirty years (Başar et al. 1984; Intriligator & Polich 1995; Roschke & Fell 1997). It has been realized that delta response dominates the P300 response (Stampfer & Başar 1985; Knyazev 2007). The equivalence has suggested a different prism to look at the late ERP components beside the averaging method. The alternative consists in focusing on the delta range of evoked cortical oscillations. The benefit of that approach is possibility to derive information from single trials (Schürmann, Başar-Eroglu, Kolev & Başar 2001). The possibility is crucial for our attempt to relate the SCR with the respective EEG epoch, i.e. to introduce the concept of the SCR-related brain oscillations.

There is also functional commonality between the late ERPs with latencies over 300 milliseconds and delta EEG activity – they are both markers of emotional processing (Klados et al. 2009; Başar, Schmiedt-Fehr, Öniz & Başar-Eroglu 2008; Güntekin & Başar 2016; Cuthbert et al. 2000; Olofsson et al. 2008; Brown et al. 2012). They are evoked by stimuli that are unexpected, infrequent, and motivationally relevant (Knyazev 2007, 2012; Cuthbert et al. 2000; Olofsson et al. 2008).

Over 50 years ago Walter Freeman (1964, 1967, 1968) proposed that the evoked oscillatory waves of the EEG are generated by multiple positive and negative forward and feedback loops. While Freeman used to rely exclusively on the within cortical neural loops in the modeling of cortical evoked potentials later authors expended the concept of neural feedback or reentrant signaling also to the neural populations out of the cerebral cortex such as thalamus (Edelman & Gally 2013).

The question of distinct effects of forward and feedback neural signals on event-related cortical responses (ERPs) has been recently posed (Woodman 2010). Both theoretical consideration (Woodman 2010) and results of computational modeling of ERP elicited by deviant stimuli (Garrido, Kilner, Kiebel & Friston 2007) have converged to the conclusion that the earliest ERP components occur due to feedforward processing of the representations of the stimuli and late ERP components with latencies longer than 200 ms are generated by backward connections. More specifically, P3a subcomponent of ERPs has been associated with the dopaminergic feedback to the cortex and P3b with the LC- noradrenergic feedback signal (Nieuwenhuis, Aston-Jones & Cohen 2005; Polich 2007).

Spectral analog of the late ERPs, delta event-related EEG oscillations, have been associated with dopamine, noradrenaline, and acetylcholine neurotransmission (Fingelkurts & Fingelkurts 2010; Başar & Düzgün 2016a). Knyazev (2007, 2012) has also pointed to the subcortical generation of both P300 and delta brain activity, mostly through dopaminergic neurotransmission. Knyazev (2007, p. 381) suggested that “a phasic omission rather than a splash of dopamine neurons firing could be conducive to the P300 [and delta activity] generation”.

### Monoaminergic signaling visible through computational modeling of the SCR

In our computational approach to the SCR we consider the SCR as an output of a series of integrations in both neurophysiological and computational meaning of the word (Branković 2011, 2012). We approximate the arousal process by a series of integrations with the integration step of 100 ms accompanied with three feedback loops. The SCR system in this perspective is viewed as a linear neurochemical oscillator suitable for mathematical modeling and estimation of the inherent control process.

We have also argued that peripheral processes, i.e. the biophysical properties of the sweat glands, do not significantly contribute to the overall dynamics of the system (Branković 2011), what is congruent with a recent view on the mechanism of the SCR (Dementienko et al. 2000). Similar assumption has been exploited in the pupil dilation model by Hoeks & Ellenbroek (1993). To paraphrase these authors: the convolution of the signal entering the sweat gland does not change the overall impulse response dramatically.

Two set of parameters have emerged through this approach – one wich refers to the hidden neural input to the SCR system and the other which characterizes the regulatory (control system) aspects of the SCR process.

We have suggested that the SCR signal and its derivatives reflect the neural signals at different integration levels in the central regulation of the arousal process. Although the nodes in our dynamic model could be realized as principal points on the path of integration and feedback regulation of the central neural signal conveying the information for emotional sweating we did not suggest that the SCR signal which we measured at the skin surface and which figures as the slow positive feedback loop in the SCR model was actually transmitted back to the brain through some receptors and ascending neural pathways and took part in the regulation of arousal. Rather, we supposed that in the central regulation of the arousal appears some neural signal (we assumed the brainstem monoaminergic activity), with similar and coherent temporal characteristics as the measured SCR, which is the real slow positive feedback signal in the control process of the SCR. We hold the similar view on the fast positive feedback signal and the negative feedback signal in our SCR model. Mathematically they appear as the first and the second derivative of the measured SCR signal but neurobiologically they correspond to two earlier nodes in the neural integration chain during the SCR process.

“Psychopharmacological dissection” of these signals through a pharmacological intervention study (Branković 2012) suggested their neurochemical substrate. The second derivative of the SCR have been associated with the phasic dopaminergic activation of the amygdala, the first derivative of the SCR with the phasic LC-noradrenergic feedback inhibition to the amygdala, and the SCR signal itself with the phasic serotonergic feedback activation in the arousal process.

### Hypothesis of the existence of the SCR-related brain oscillations and their feedback origin

Before we expose the hypotheses of the SCR-related brain oscillations it is pertinent to question why the SCR-related brain oscillations have not been already observed and described before. According to Vaughan (1969, p.46) “sufficiently prominent or distinctive psychological events may serve as time references for averaging, in addition to stimuli and motor responses. The term ‘event-related potentials’ (ERP) is proposed to designate the general class of potentials that display stable time relationships to a definable reference event.” Skin conductance responses and ERPs emerge on different time scales and only recent computational dealing with the SCR which identifies the beginning event, onset of the SCR process on the hundred-millisecond scale (“temporal microscope” Branković 2011, 2012) provided a definable time reference of the SCR signal which can be related to the EEG signal.

The hypothesis of the existence of the SCR-related brain oscillations is based on the following chain of reasoning. Firstly, there is functional equivalence between the late ERPs – delta oscillatory brain responses and the SCR system. Secondly, we rely on the common neurochemical feedback nature of the late ERPs – delta activity and the signals involved in the control process of the SCR. Finally, we point to spectral parallelism between the models of the late ERPs – delta oscillatory brain responses and the SCR system.

Functional equivalence between the later ERP components with latencies over 300 milliseconds (P3 and LPP), delta oscillatory responding and the SCR consists in that that they all indicate emotional processing (Klados et al. 2009; Başar et al. 2008; Güntekin & Başar 2016; Knyazev 2007, 2012; Cuthbert et al. 2000; Olofsson et al. 2008; Brown et al. 2012; Boucsein 2012). There is also evidence for positive correlation between the LPP and the SCR (Cuthbert et al. 2000).

Common, phasic monoaminergic feedback mechanisms have been implied as both the origin of the P3 wave and regulatory signals in the SCR process. According to Nieuwenhuis, De Geus & Aston-Jones (2011) P3 is a correlate of the phasic “LC-NE activity. Further, LC-NE system in the brain is a central analogue of the peripheral sympathetic nervous system [including SCR] … two systems often operate in an integrated fashion… autonomic components of the orienting response and the P3 are intimately coupled” (*ibid.* p.168). Both the neuroanatomical model of P3 component (Nieuwenhuis, De Geus & Aston-Jones 2011) and our model of the SCR system (Branković 2008) involve the projections of the hypothalamic paraventricular nucleus that inervate LC through nucleus paragigantocellularis. Both models have apostrophized feeback, regulatory function of this output arm of the hypothalamus. A difference between the models consists only in that that Nieuwenhuis, De Geus & Aston-Jones (2011) have pointed to the cortical effect of the phasic LC-NE activity (P3) and we have suggested the role of phasic LC-NE activity in the regulation of the peripheral sympathetic responding (the SCR) through the feedback inhibition of the amygdala (Brankovic 2008). The difference is illustrative and we hypothesize here that it holds also for other brainstem monoamines. For instance, Polich (2007) has associated dopamine and P300b wave.

While our computational modeling of the SCR (Branković 2012) enables estimation of feedback dopaminergic, noradrenergic, and serotonergic effects at the level of the amygdala, we are also interested in cortical electrophysiological effects of dopamine, noradrenaline, and serotonin (Figure 1). These neurotransmitter systems have been recently conceptualized as “the Modulatory network” of the brainstem involved in emotion processing (Venkatraman, Edlow & Immordino-Yang 2017).

There is also spectral parallelism between the late ERPs-delta activity and the signals of the SCR system. Delta frequency range dominates the P300 and LPP response (Stampfer & Başar 1985; Knyazev 2007). On the other hand, signals involved in the SCR process (the first three derivatives and the SCR signal itself) are in the range 0.01-5 Hz (Branković 2011). That is why we expect that if there were such a thing as the SCR-related brain oscillations, it would be a delta phenomenon and the search for it should be focused on the delta frequency range of the EEG.

In other words, since evidence points to functional, neurochemical, and spectral parallelism between the late ERPs – delta oscillatory brain responses and the SCR system we assume that there is delta EEG activity which could be related to the skin conductance responding. That delta activity we designate as the SCR-related oscillations. One more assumption should be included in these reasoning. There is a time-shift between the brain process (and simultaneous EEG signal) and the recorded SCR signal due to slow conduction velocity through peripheral sympathetic unmyelinated C fibers. Since latency period for the SCR onset after a stimulus is assumed to be 1-3.5 seconds (cf. Staib et al. 2015) we expect that the time-lag between the SCR and the respective delta SCR-related EEG oscillations would be 0.5-3 seconds.

Bringing together our theorizing on the monoaminergic nature of the feedback regulatory signals in the SCR system (Branković 2012) with the hypothesis of the brainstem monoaminergic origin of the late ERPs-delta activity (Nieuwenhuis, Aston-Jones & Cohen 2005; Polich 2007; Knyazev 2007; Fingelkurts & Fingelkurts 2010) we come to a theoretical possibility that there is a connectivity between the SCR and its derivatives (reflecting the brainstem monoaminergic signaling) as pridictor signals and the respective, predected, delta EEG epoch – the assumed SCR-related oscillations (Figure 1).

## Participants and Methods

### Participants

Twenty six healthy volunteers with age range from 20 to 44, with no history of psychiatric and neurological treatment took part in the study. The participants were postgraduate and graduate students of humanities and medicine. Three participants (2 men and 1 woman) appeared to be non-responders for the SCR. Of the twenty three SCR responders 13 were men and 10 women with an average of 15.5 years of education. All the subjects were right-handed. They possessed corrected to normal to normal vision. All participants were briefed about the experiment. All of them showed their consent and signed the consent form before the recording session. The Human Research Ethics Committee of the University of Belgrade approved the study protocol.

### Experimental procedure

Eleven short stories from the contemporary literature (without erotic and aversive content) have been chosen to elicit pleasant excitement in subjects. The stories were divided into meaningful fragments and presented to the subjects as slide presentations on a 15″ monitor. The stimulus material and experimental procedure have been explained in Branković (2008, 2011). The subjects were instructed to read the stories as they usually do during leisure time and to switch to the next slide by their own clicking a computer mouse. During the slide-presentation EEG, skin conductance, heart rate, and respiration of subjects were recorded.

The slide-presentation and psychophysiological recording started after a five minutes adaptation period to experimental room and equipment. All subjects were tested during afternoons (2-8 P.M.). The room temperature was held between 20 and 22°C.

### Data acquisition

The psychophysiological measurement has been performed using the PowerLab^®^ 4/25, signal conditioners Bio Amp, GSR Amp, and the software for digital data acquisition LabChart^®^ 7 for Windows^®^ with a sampling rate of 100 Hz.

The subjects washed their hands with soap and warm water before the montage of electrodes for skin conductance on the middle phalanges of the second and fourth finger on the left hand.

We used only three midline channels to record the EEG signal (F_pz_, F_z_, and P_z_ according to the 10-20 International System). F_pz_ was used as a ground, F_z_ as a reference, and P_z_ as active electrode. The selection was based on the functional differentiation of the brain: F_z_ is located near motivational centers and the signal from P_z_ is associated with activity of perception and differentiation (Teplan 2002). These sites also show the largest amplitudes of P3 (Polich 2007; Nieuwenhuis, Aston-Jones G & Cohen 2005; Nieuwenhuis, De Geus & Aston-Jones 2011) and LPP (de Rover, Brown, Boot, Hajcak, van Noorden, van der Wee & Nieuwenhuis 2012; Cuthbert, Schupp, Bradly, Birbaumer & Lang 2000).

Epochs contaminated by eye and other artifacts were manually rejected offline. Delta EEG component defined as the 1-4 Hz band has been derived offline in LabChart^®^ 7 for Windows^®^ which implements linear phase finite impulse response filters designed using the window method with a Kaiser window (beta=4.86), giving pass and stop band ripple of less than 0.5%.

### Identification of the time delay between the EEG and the SCR signal

Relying on our hypotheses about (1) the SCR-related oscillations as a delta process and (2) the connectivity between the delta EEG and the SCR system signals, and taking into consideration a variable latency period for the SCR onset we developed a method for detecting the delta EEG segments which correspond to the putative SCR-related oscillations. The method consists in finding firstly the three candidates for the delta EEG epochs and choosing then the one which maximizes the regression coefficient. Experimenting with different lengths of the epochs the duration of two seconds appeared optimal for the purpose.

The three candidates for the SCR-related EEG epochs are looked for on the delta EEG signal with the beginning located in the interval from 3 seconds up to 0.5 seconds before the onset of the SCR (defined as the maximum of its third derivative, *see* Branković 2011). There are thee kinds of these two second candidate epochs:

1. the delta EEG segment which starts at the point of the maximum slope of the delta EEG signal in the searched interval (presuming to reflect the onset of a new forcing input to the oscillatory system);
2. the delta EEG segment which corresponds to the maximum cross-correlation between the segment and the third derivative of the SCR (a classical technique for time delay estimation);
3. the delta EEG segment which corresponds to the maximum sum of the correlation coefficients which refer to the cross-correlations between the segment and all first three derivatives of the SCR and the SCR signal itself.

### Multiple time series regression

The thesis on the connectivity between the SCR system signals and the delta EEG component was operationalized through a simple model of effective connectivity – a multiple time series regression model (Büchel & Friston 2002; Lieder, Stephan, Daunizeau, Garrido & Friston 2013). Time delay between the SCR system signals at adjacent integration levels have been varied from 0 to 200 ms and the values which maximized the regression coefficient have been selected.

### Monte Carlo test of significance of the time series regression

Statistical significance of time series regressions was assessed using a Monte Carlo approach, i.e. calculating the empirical p-value of the null hypothesis. We tested for each individual participant the significance of difference between (1) her/ his median value of the explanatory powers of regressions (R^2^) obtained on true data (the two-second SCR system signals and the SCR-related delta-EEG oscillations) and (2) the median of the R^2^ obtained on 500 random fake delta-EEG two-second samples which were unrelated to the SCRs.

Unrelatedness with the SCR has been provided by the condition in the randomization that the beginning of the EEG epoch does not precede less than six seconds any SCR onset. The time delay between the EEG and the SCR signal, and identification of the SCR‴ segment beginning and lags of other SCR system signals in the regression analyses have been performed in the same way for the fake samples as it had been done for the true data.

## Results

### The SCR-related brain oscillations

Grand average of the SCR onset-locked unfiltered EEG oscillations baselined at the interval 3400-3200 ms before the SCR onset is presented on the Figure 2A. There is no marked regularity which could be seen except the initial positive deflection about 3 s before the SCR onset.

**Figure 2.**
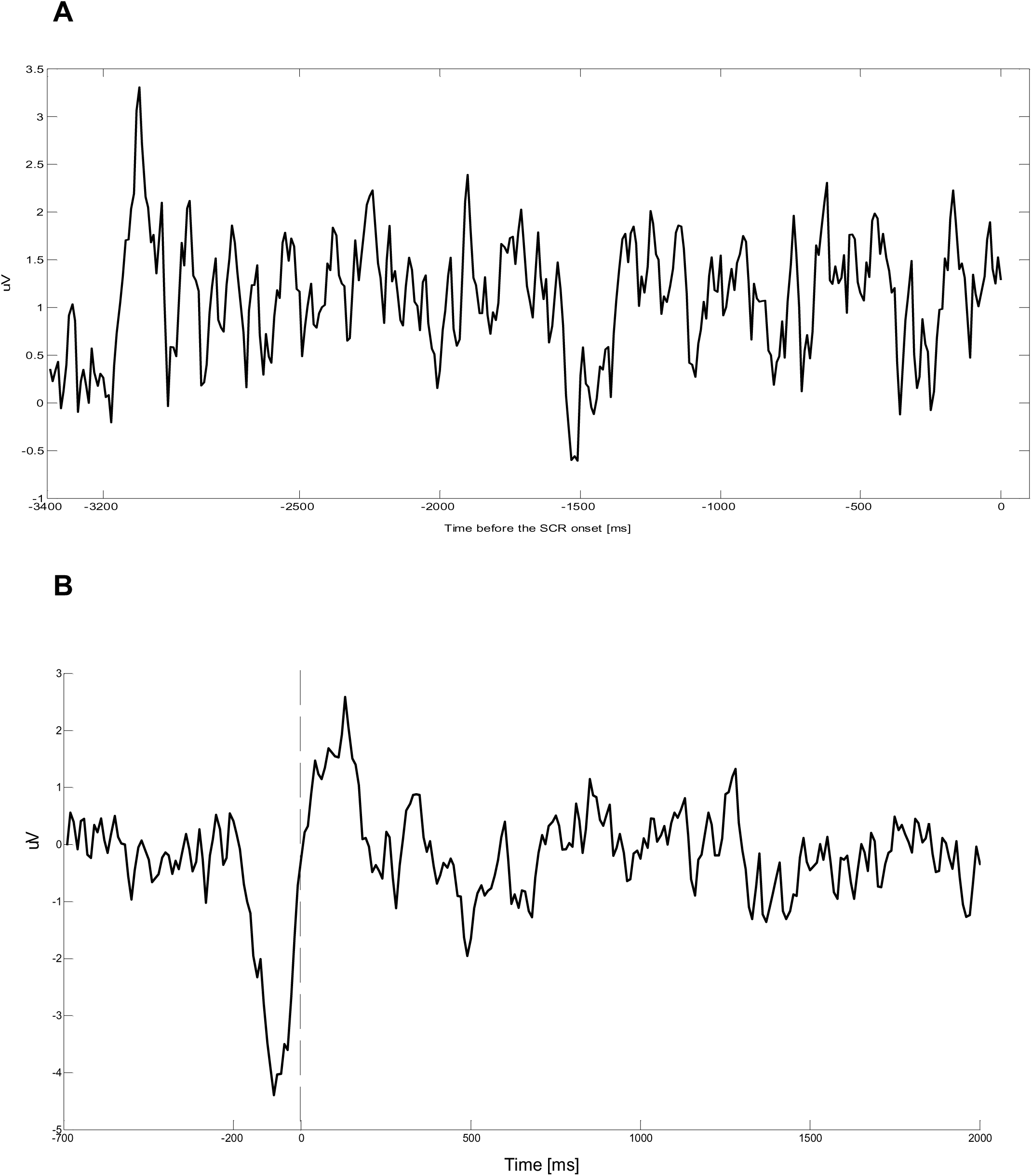
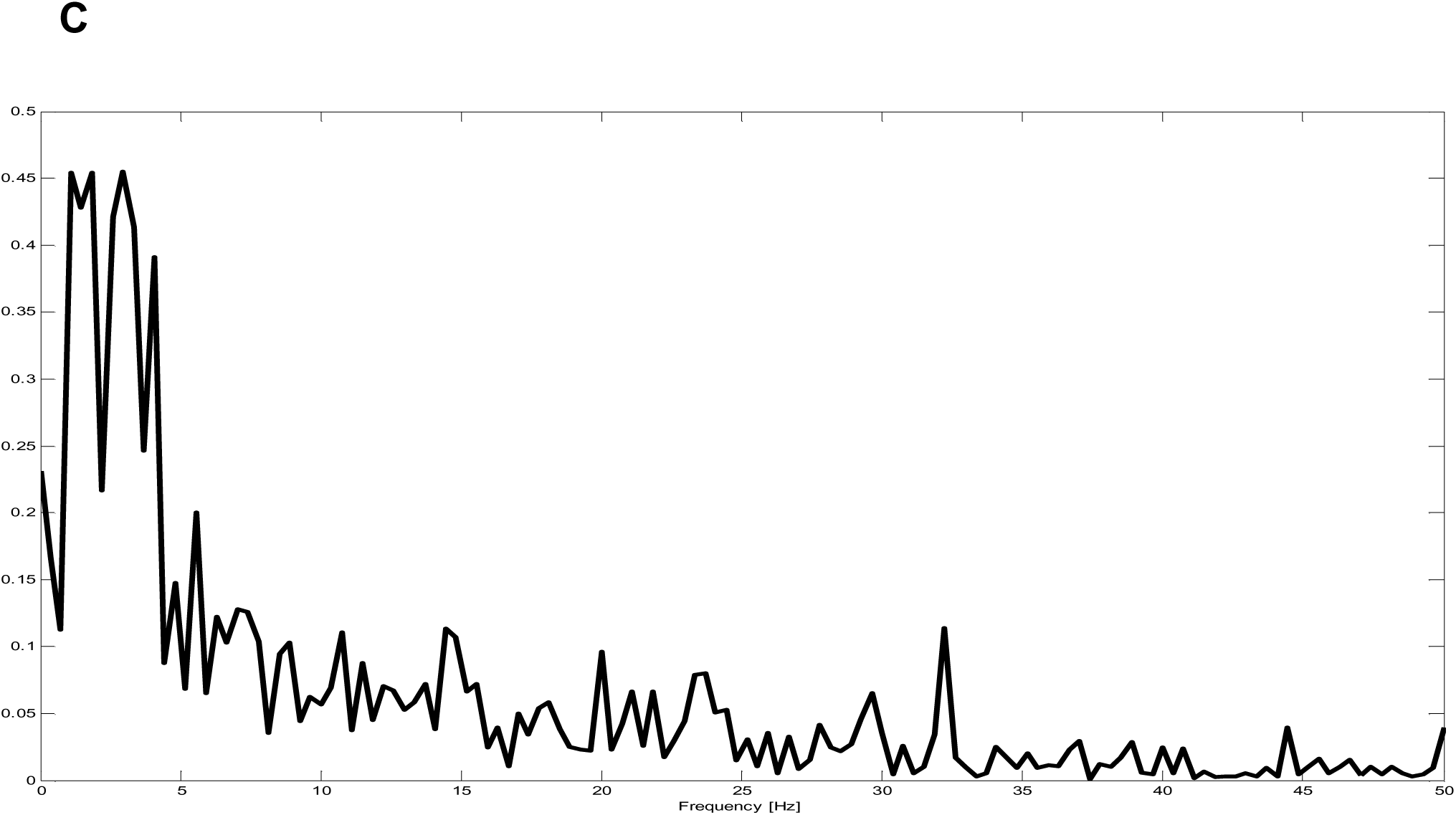
Grand average of the SCR onset-locked unfiltered EEG oscillations **(A)**, grand average of the identified (data derived) unfiltered SCR-related EEG oscillations **(B)**, and its amplitude spectrum **(C)**.

On the other hand, the grand average of the unfiltered SCR related two second EEG epochs with beginnings determined through the method explained in *Participants and Methods (Identification of the time delay between the EEG and the SCR signal)* is more revealing (Figure 2B). Its amplitude spectrum points to dominant delta activity, to a lesser extent to theta activity, and damping of the higher frequencies through the averaging procedure (Figure 2C). In that way, the hypothesis about the existence of the SCR-related oscillations and their delta nature is confirmed.

An unexpected finding here is that the SCR-related oscillations are more complex than we postulated. In addition to the two second EEG segment which is correlated with the SCR system signals they comprise an initial segment of 200 ms – a deep negative deflection. This negative wave peaks at the middle of the initial segment, about 100 ms after the onset of the evoked oscillatory response and 100 ms before the start of the two second epoch which is influenced by the SCR system signals (*see also* Figure 2A).

To conclude, the SCR-related brain oscillations are composed of two adjacent EEG segments. The first one is a negative deflection with duration of 200 ms and it is uncorrelated with the SCR system signals. The second part of the SCR-related oscillations is the postulated two second EEG segment which is correlated with the SCR system signals.

### The SCR-related delta brain oscillations as superposition of the SCR, its derivatives, and their interactions

The hypothesis about the connectivity between the SCR and its derivatives as predictor signals and the respective SCR-related brain delta oscillations as an observed response has been operationalized through a multiple time series regression model. The response variable is the two second SCR-related delta EEG epoch with beginning determined through the method explained in *Participants and Methods (Identification of the time delay between the EEG and the SCR signal)*. Experimenting with different predictor sets we have chosen the following 16 predictor variables: the SCR, its first three derivatives, their 11 possible mutual interactions, and the “hidden input” to the SCR system (determined as explained in Branković 2011). The sample size in the regression has been 200 (two second signal epochs with sampling rates of 100 Hz).

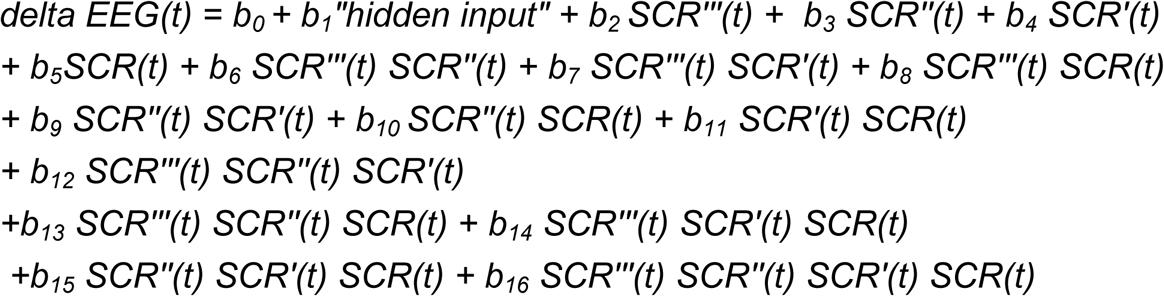

An example of the modeling of the two second EEG segment through the SCR system signals is presented on the Figure 3. The distribution of the method of determination of the beginning of the EEG segment which yielded to the most successful regression model is the following: (1) the point of the maximum slope method was the choice in 109 cases of the SCR-related epochs; (2) the maximum cross-correlation between the EEG segment and the third derivative of the SCR was the choice in 180 cases; and (3) the maximum sum of the cross-correlation coefficients was the choice in 182 cases. Average number of the identified SCR-related delta EEG epochs per subject in our sample was 20 (range: 5-57).

**Figure 3.**
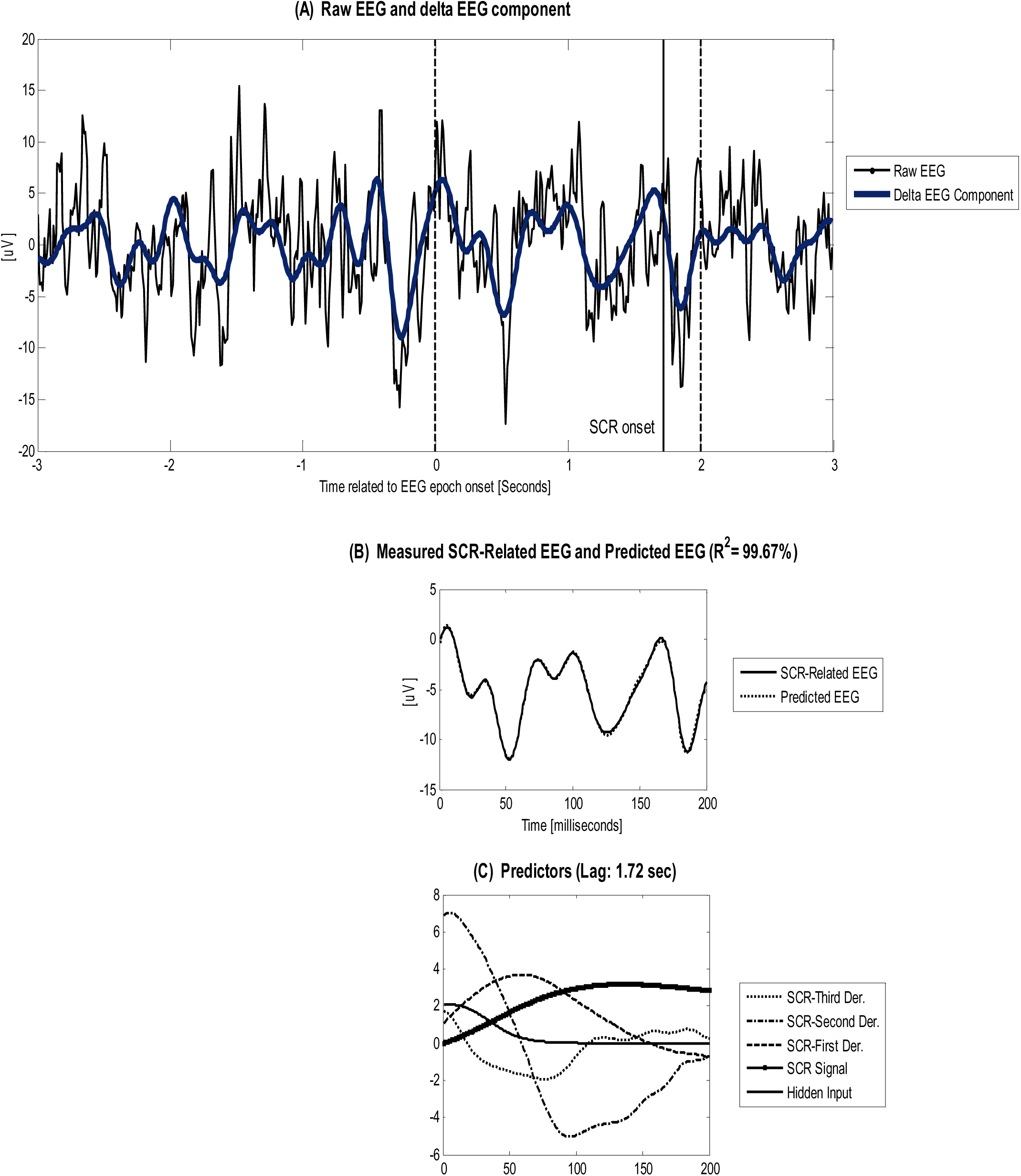
An example of the modeling of delta SCR-related oscillations by regression to the SCR system signals.

The distributions of the advance of the SCR related epochs of the EEG signal in respect to the SCR onset (identified by the third derivative of the SCR) and time lags of the individual SCR system signals are presented on the Figure 4. The time correction for the SCR‴ as a predictor signal refers to the SCR‴ signal itself. Lags of the other SCR system signals refer to the signal at the lower adjacent integration level, i.e. the lag of the SCR″ refers to the SCR‴ signal, the lag of the SCR’ refers to the SCR″ signal, and the lag of the SCR refers to the SCR’ signal.

**Figure 4.**
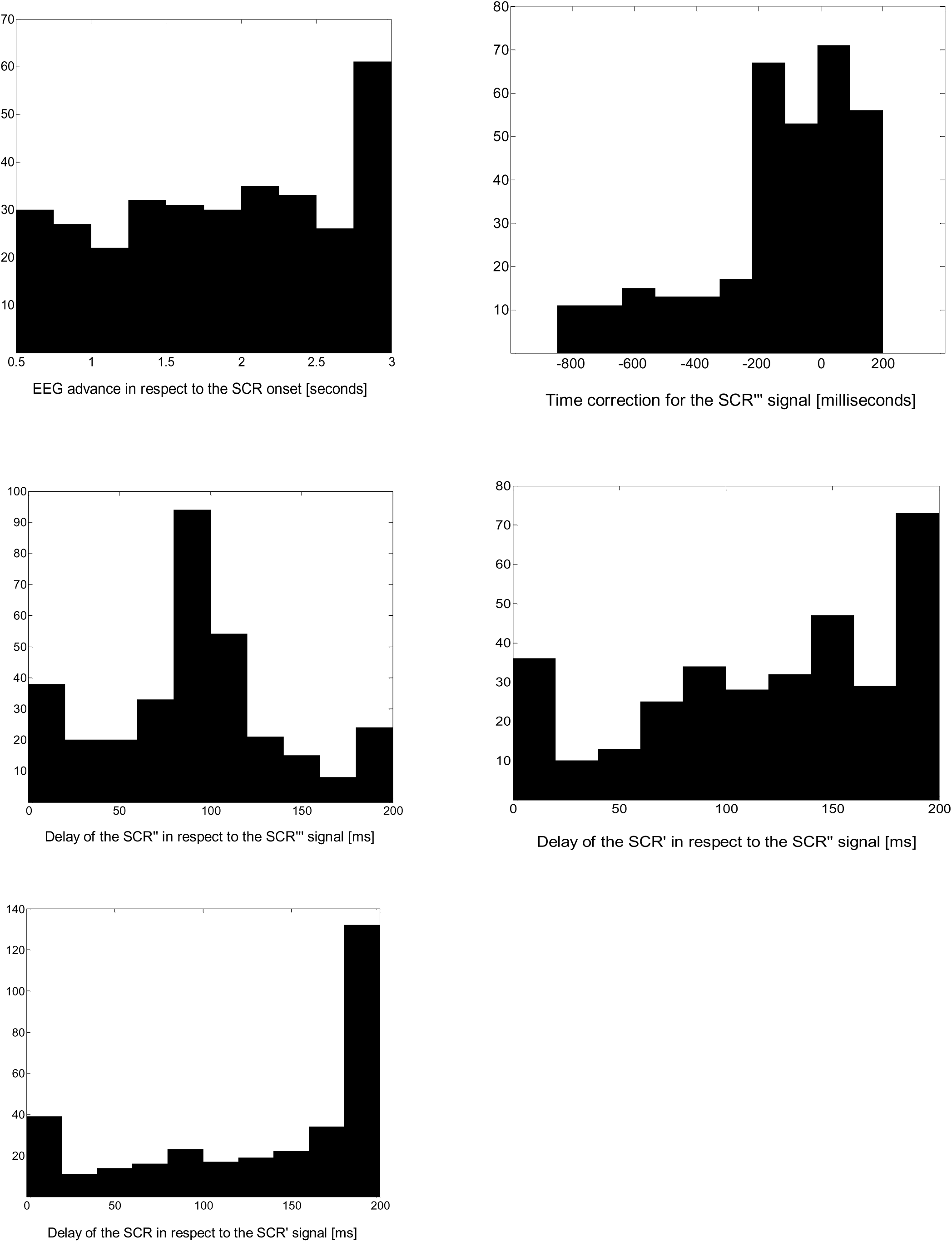
The time delay metrics (histograms of the lag values among the signals).

The adjusted R^2^ values in the whole sample of regression models are in the range 0.8165-0.9998, with the median of 0.9832. The predictor set showed significance at the level p<0.001 for all estimated coefficients except in sporadic cases and F-statistics showed significance p<0.001 for all regression models.

Regression coefficients manifested temporal fluctuations throughout the recording session (Figure 5) pointing to dynamic changes in effective connectivity.

**Figure 5.**
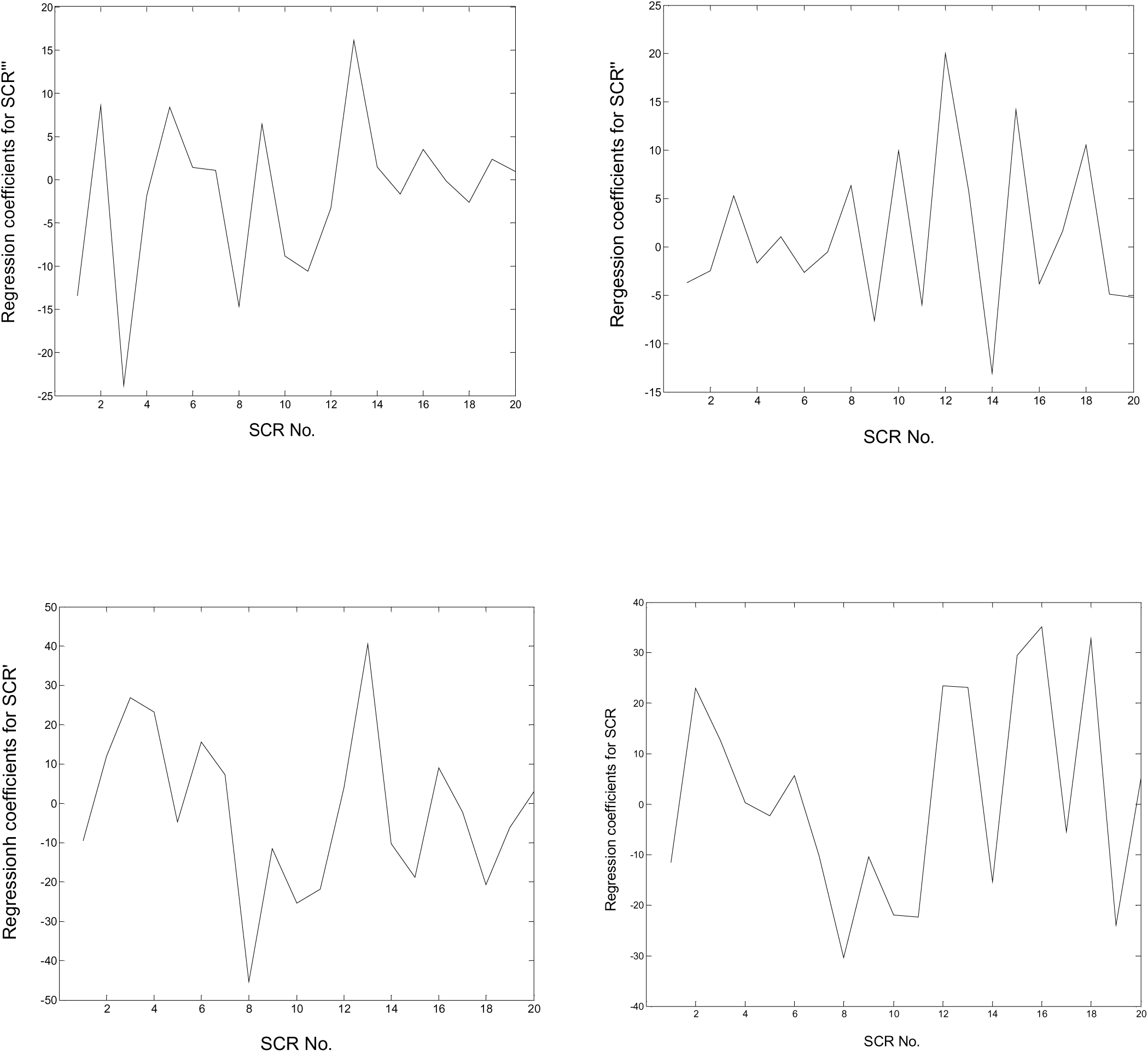
Temporal fluctuations of individual regression parameters (participant No.14): dynamic effective connectivity.

### Monte Carlo evidence for the causal link between the SCR signal and its derivatives, and related delta EEG oscillations

In order to test causal nature of the obtained regression model between the SCR system signals and the respective delta EEG oscillations we performed a Monte Carlo analysis. The regression model with full set of 16 predictors did not show significance of difference in explanatory power of regression (R^2^) between the true, SCR-related delta EEG oscillations, and fake EEG samples which are unrelated to the SCR. Taking into consideration the high values of R^2^ in both true and fake samples we have assumed that the failure of obtaining statistical significance could be caused by the lack of discriminating power of the full set of predictors for the purpose of the Monte Carlo analysis, i.e. by the ceiling effect.

In order to check this possibility we have performed Monte Carlo analyses of the regression models with reduced numbers of predictor variables. Three levels of simplified regression models have been applied: the model with 13 predictors (the first 13 predictor variables of the full set of 16 predictors), the model with 9 predictors (the first 9 predictors of the full set), and the model with 5 predictors (the first 5 predictors of the full set). Through the simplification of the regression model we obtained significance of difference in R^2^ between the true and fake samples, in the sense of significantly greater median of R^2^ values in the true samples at the level of p<0.005 in 13 participants (Figure 6).

**Figure 6.**
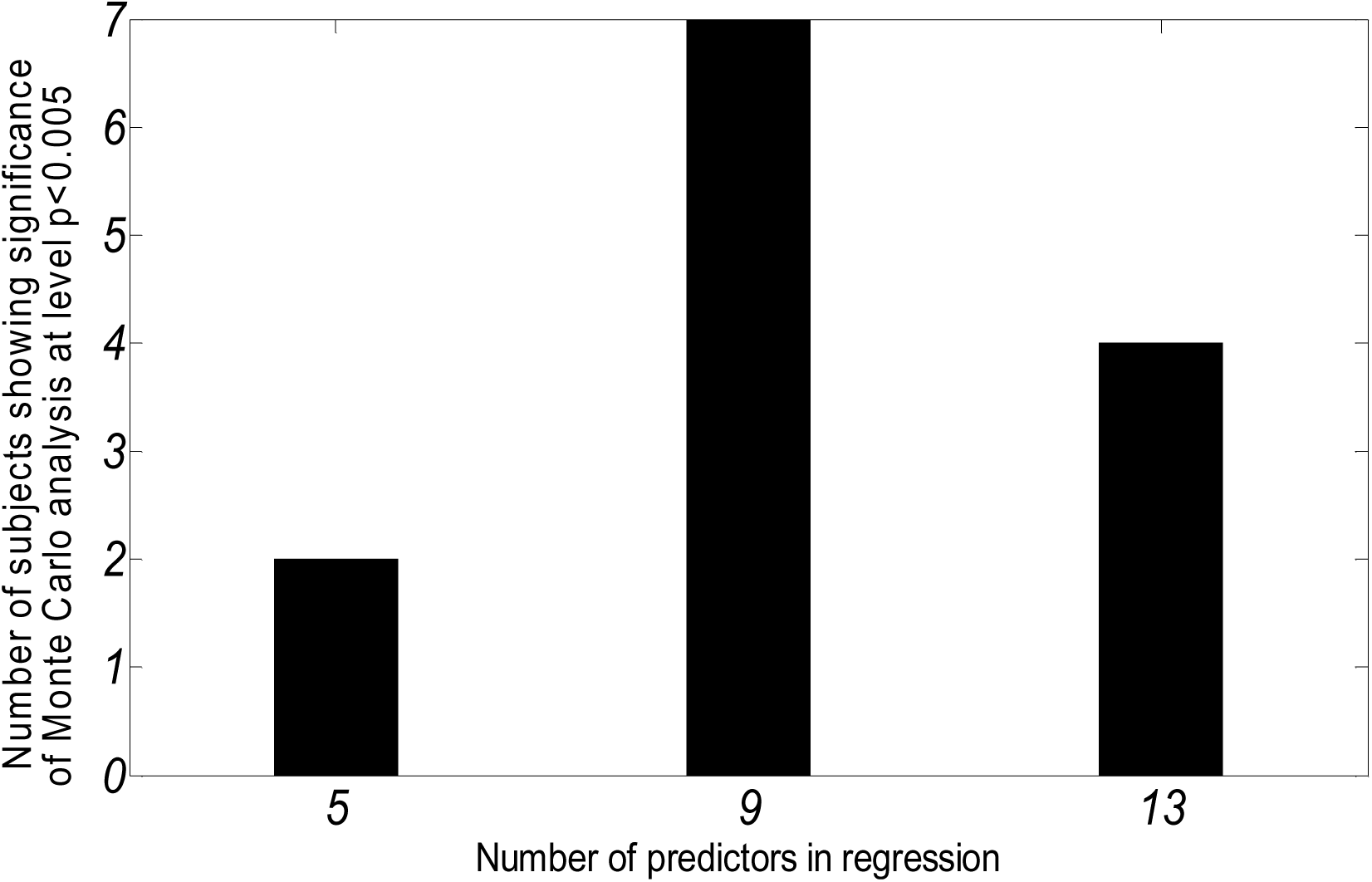
Histogram of the number of predictors in regression model which yielded to significance in Monte Carlo testing of the EEG to SCR system signals regression.

## Discussion

In contrast to the classical approach to event-related potentials and oscillations which relies on either stimulus- or response-locked time reference we have introduced here an approach which we could assign as “oscillatory process – related oscillations”. We have found that the grand average of the SCR onset-locked unfiltered EEG oscillations is blurred by variable latency period of the SCR. On the other hand, the new approach reveals more information by collating the EEG epochs based on data derived information about the oscillatory process. The method enables us to overcome the variability of sympathetic nerve conduction velocity, i.e. the variability of latency period of the SCR. The variability used to blur the information on the oscillatory brain process related to the SCR which is available on the EEG signal.

In agreement with our hypothesis the SCR-related brain oscillations have been identified as delta frequency process. Our approach to identify the SCR-related oscillations through correlations with the SCR system signals (Branković 2011) led not only to confirmation of the hypothesis but also revealed that the phenomenon of the SCR-related oscillations is more complex than we presumed and broader than the effects of the SCR system. It comprises in addition an initial 200 ms EEG segment, a deep negative deflection, which is unlinked with the SCR system. Due to (1) its temporal distance from the baseline activity (i.e. a latency of 100-200 ms) and (2) character of the evoking situation for the emotional SCR (Branković 2011) which implies unfulfillment of expectations in emotional communication (Branković 2001) it is tempting to question weather this negative wave is somehow related to the mismatch negativity (Lieder, Stephan, Daunizeau, Garrido & Friston 2013).

We have expected that delays between the SCR-related brain oscillations and the respective SCR would be 0.5-3 seconds. Variability of the delays could be explained by the large temporal dispersion of the slow, unmyelinated, peripheral, sympathetic C fibers (Hari & Parkkonen 2015). The delays have the mode value at the upper limit of the chosen interval suggesting that even longer delay periods are actually involved.

The SCR-related brain oscillations are here modeled as the superposition of the SCR, its derivatives, and their interactions. The regression analysis showed that the predictor set explains a high majority of variance of the delta EEG fluctuations related to the SCR. Relying on the Monte Carlo evidence for the causal link between the SCR system and the related delta EEG we conclude that our hypothesis about the connectivity between the SCR system signals as predictor signals and the respective delta brain oscillations as an response is confirmed.

### Significance of the finding and explanation of the causal link between the SCR system signals and the related delta EEG

There are several aspects of the significance of the Monte Carlo evidence for the causal link between the signals of the SCR system (Branković 2008, 2011) and the related delta EEG fluctuations. We firstly point to a significance which is independent of the theoretical background which led to the present hypothesis and of the explanation of the neurobiological nature of the link which we propose further. Namely, the finding points to existence of a mechanism through which the SCR system and EEG fluctuations are connected. In that way, the finding justifies the central nervous system interpretation of the dynamic models of the SCR (Branković 2008, 2011, 2012).

Another aspect of the significance of the finding is that it speaks in favor both of the models of the monoaminergic origin of the late ERPs-delta activity (Nieuwenhuis, Aston-Jones & Cohen 2005; Polich 2007; Fingelkurts & Fingelkurts 2010) and of the monoaminergic nature of the SCR system signals (Branković 2012). Since the implication derived from the synthesis of these two models (the hypothesis of connectivity between the SCR system signals and the related delta EEG) is confirmed, we can conclude that these models are mutually congruent and thereby justified in the present study. In the light of these models, the evidence about the causal link between the SCR system signals and the related delta EEG segment points to the monoaminergic signaling as the neural mechanism of this connectivity. An interpretation of this theoretical synthesis is that in contrast to the “one ERP – one neuromodulator” approach (Nieuwenhuis, Aston-Jones & Cohen 2005; Polich 2007) we propose here an “effective connectivity” approach to the SCR-related brain oscillations.

According to this integrated model the regression coefficients in the performed time series regression reflect the cortical phasic effects of the brainstem monoamines, the amygdala, and their interactions. While the feedback loops’ gains in the SCR system have been interpreted as measures of the phasic brainstem monoaminergic effects on the amygdala-hippocampus circuit (Branković 2012), the regression coefficients in the present regression model correspond to the respective cortical components and their mutual interactions.

The significance of the regression coefficients associated with the interaction factors in the regression model could be understood in a way that both the monoaminergic interactions and also brainstem – amygdala interactions contribute to the resulting cortical superposition of the phasic monoaminergic and amygdala signaling. In favor of that explanation speaks the evidence about the adrenergic beta-receptor-mediated modulation of the late positive potential (LPP) through activation of the amygdala (de Rover, Brown, Boot, Hajcak et. al. 2012).

Our setting the algorithm with the limit of two hundred milliseconds allowed time lag between signals from adjacent integration level of the SCR system (i.e., the third, second, first derivative, and the SCR signal itself) enabled a successful regression of the SCR-related brain oscillations. On the other hand, the chosen setting is neurophysiologically realistic and corresponds to the identified dynamics and time delays of dopamine, noradrenaline, and serotonin release in targeting areas (Hashemi, Dankoski, Lama, Wood, Takmakov & Wightman 2012; Muller, Joseph, Slesinger & Kleinfeld 2014).

The finding of temporal fluctuations of the regression parameters in our model of the causal link between the SCR system signals and the related delta EEG points to oscillations of effective connectivity, i.e. the dynamic connectivity. The phenomenon is well documented and seems promising for “extracting meaningful information from functional neuroimaging data” (Breakspear 2004, p.205). It could be valuable to identify distinct patterns of the parameter variation in our connectivity model for specific diagnostic entities (cf. Kaiser, Whitfield-Gabrieli, Dillon et al. 2015; Rashid, Damaraju, Pearlson & Calhoun 2014; Jia, Hu & Deshpande 2014) and to analyze the process of parameter variation itself. An open question is what is the neural origin, mechanism, of dynamic functional connectivity and the role of the brainstem and subcortical structures in its regulation has been already suggested (Hutchison et al. 2013).

Our interpretation of the causal link between the SCR system signals and the related delta EEG is in accordance with the thinking of Makeig & Onton (2009) that trial-to-trial EEG variability reflects not “ERP noise” but instead the brain’s carefully constructed response to the highly individual stimulus- and context-defined unfolding situation. Moreover, “changing EEG dynamics index changes in and between local synchronies that are driven or affected by a variety of mechanisms including sensory information as well as broadly projecting brainstem-based arousal or ‘value’ systems identified by their central neurotransmitters – dopamine, acetylcholine, serotonin, norepinephrine, etc.” Makeig & Onton (2009). If this is the case, future research could show how the regression coefficients in our regression model relate to diagnosis and treatment of certain mental disorders.

The Monte Carlo test of the causal link between the SCR system’s signals and the related delta EEG did not demonstrate significance in 43% of the participants. Rather than concluding that the causal link is a feature of some participants and not of the others, we assume the following explanations for this inconsistency of the results. We distinguish principally two kinds of the possible reasons for the inconsistency – methodological and fundamental. The methodological circumstances could prevent reaching the statistical significance of the Monte Carlo testing through the lack of power of the applied regression predictor sets to discriminate between true and fake data, i.e. through the floor and ceiling effect due to similar values of R^2^ in the regression models. Actually, in more than a half of the participants we confirmed the ceiling effect – reduction in the number of predictors, i.e. elimination of the interactive factors in the regression model, brought to the significance of the difference between the true and fake samples. Another methodological factor which appeared effective in revealing the causality between the SCR system and the EEG is the treatment of the time delays between the signals. A trial with alteration of allowed values for the time lags in the algorithm revealed significance in the Monte Carlo testing in one participant.

The fundamental reason for not reaching the significance of difference between the true and fake samples in the Monte Carlo testing of the causal link between the SCR system and the EEG could be the known segmental structure, uniformity and repetitive nature of the EEG (Kaplan 1999; Fingelkurts & Fingelkurts 2015). There are segments of the EEG lasting up to several seconds, separated by abruptly jumps, which mark the discontinuities of relatively stable functioning (i.e. free of forcing inputs) of the local neuronal networks. It has been proposed that during these stationary periods a particular brain system executes separate operations. It has been shown that different frequency EEG bands are featured with different segmental structure. The length of segments is from hundreds milliseconds up to several seconds and longer for delta activity than for higher frequency bands but do not exceeds 4 seconds (Elul 1969; Inouye, Toi & Matsumoto 1995; Kaplan 1999). It is possible that at rest, during the SCR-free periods, the brain (cortical) activity reflects spontaneous events (Petridou, Gaudes, Dryden, Francis & Gowland 2012; Tagliazucchi, Balenzuela, Fraiman & Chialvo 2012) which include brainstem monoaminergic modulation similar to that involved in the SCR regulation, but without an accompanied peripheral sympathetic output – the SCR.

### Embedding the Skin Conductance Response System into the “Brain-Body Dynamic Syncytium”

Our thesis about the SCR-related delta brain oscillations as a weighted sum of the SCR signal, its derivatives, and their interactions could be regarded from the standpoint of a recently emerged more general model – the brain-body dynamic syncytium” hypothesis (Başar 2012). According to that model the body and the brain use the same transmitters and the same frequences for tuning of the brain-body interaction. Başar has pointed to “the multiple oscillatory processes… between brain and body visceral organs and tissues within a stuctural and functional continuum termed ‘Brain-Body-Mind functional syncytium’. Within such ‘functional syncytium’, quasi-invariant nervous oscillatory processes at different frequences would ensure a stochastic unstable transfer of (excitatory and inhibitory) signals among different brain nodes by means of neurotransmitter systems” (*ibid.*, p.286). Further, “Oscillations and neurotransmitters work together to form one combined activity. Therefore, the web of ‘*oscillations and neurotransmitters*’ can also be considered as building blocks for function. Spontaneous and event-related oscillations in the CNS and vegetative organs are all embedded in biochemical pathways (neurotransmitters). These oscillatory processes can be considered as manifestations and building units for brain-body functioning” (Başar & Düzgün 2016b, p. 201).

The Başar’s theory implicitly suggests that peripheral oscillations of the vegetative organs (e.g. the SCR) convey information about the brain since they are connected through a dynamic syncytium. The model point to a possibility that we could define a broader connectivity framework which includes also peripheral psychophysiological measures in addition to the central, at the brain recorded signals (e.g. EEG and fMRI). Moreover, it could be the case that this broader, the “brain-body connectivity framework", is more suitable, more revealing connectivity projection for the assessment of the brainstem monoaminergic signaling than connectivity analysis limitted to signals which are all not only generated but also recorded at the brain itself. The finding of the present study about the causal link between the SCR signal and its derivatives on one side with the related delta EEG epoch on the other side lends support to the thesis that skin (the body) and the brain are more oscillatory connected than we used to think (Başar 2012).

## References

Başar E. (2012): Is research on brain oscillations in a new “take off – state” in integrative brain function. International Journal of Psychophysiology 85: 285–290. doi: 10.1016/j.ijpsycho.2012.07.180

Başar E., Başar –Erogly C., Rosen B. et al. (1984): A new approach to endogenous event-related potentials in man: relation between EEG and P300 wave. International Journal of Neuroscience 24: 161–180.

Başar E. & Düzgün A. (2016a): How the brain is working? Research on brain oscillations and connectivities in a new “Take-Off state”. International Journal of Psychophysiology 103: 3–11.

Başar E. & Düzgün A. (2016b): The brain as a working syncytium and memory as a continuum in a hyper time space: Oscillations lead to a new model. International Journal of Psychophysiology 103: 199–214.

Başar E. & Güntekin B. (2008): A review of brain oscillations in cognitive disorders and the role of neurotransmitters. Brain Research 1235: 172–193.

Başar E., Schmiedt-Fehr C., Öniz A. & Başar-Eroğlu C. (2008): Brain oscillations evoked by the face of a loved person. Brain Research 1214: 105–115.

Boucsein W. (2012): Electrodermal Activity (2^nd^ ed.), New York: Springer.

Branković S. (2001): What have the sexual, the attachment, and the exploratory motivation in common?: The theory of informational needs. Psychiatr Danub 13: 31–34.

Branković S.B. (2008): System identification of skin conductance response in depression – an attempt to probe the neurochemistry of limbic system. Psychiatria Danubina 20: 296–308.

Branković S. (2011): Interlinked positive and negative feedback loops design emotional sweating. Psychiatr Danub 23:10–20.

Branković S. (2012): Assessment of brain monoaminergic signaling through mathematical modeling of skin conductance response. In: C.M. Contreras (Ed.): Neuroscience – Dealing with Frontiers, 83-108. InTech, Rijeka, Croatia.

Branković S. (2014): Skin conductance response (SCR) related brain potentials – a new approach to assess the brainstem monoaminergic signaling. In: Abstract Book of the DGPPN Congress 2014. German Association for Psychiatry and Psychotherapy, Berlin, 2014.

Breakspear M. (2004): “Dynamic” connectivity in neural systems: theoretical and empirical considerations. Neuroinformatics 2: 205–226.

Brown S.B.R.E., van Steenbergen H., Band G.P.H., de Rover M. & Nieuwenhuis S. (2012): Functional significance of the emotion-related late positive potential. Front. Hum. Neurosci. 6: 33. doi: 10.3389/fnhum.2012.00033

Büchel C., Friston K. (2002): Interactions among neuronal systems assessed with functional neuroimaging. In: K.L. Davis, D. Charny, J.T. Coyle & C. Nemeroff (Eds.): Neuropsychopharmacology: The Fifth Generation of Progress, 383–392. American College of Neuropsychopharmacology.

Cuthbert B.N., Schupp H.T., Bradly M.M., Birbaumer N. & Lang P.J. (2000): Brain potentials in affective picture processing: covariation with autonomic arousal and affective report. Biological psychology 52: 95–111.

Dementienko V.V., Dorokhov V.B., Koreneva L.G., Markov A.G., Tarasov A.V., Shakhnarovich V.M. (2000): The hypothesis of the nature of electrodermal reactions. Human Physiol. 26: 232–239.

de Rover M., Brown S.B.R.E., Boot N., Hajcak G., van Noorden M.S., van der Wee N.J.A. & Nieuwenhuis S. (2012): Beta receptor-mediated modulation of the late positive potential in humans. Psychopharmacology 219: 971–979.

Edelman G.E. & Gally J.A. (2013): Reentry: a key mechanism for integration of brain function. Frontiers in Integrative Neuroscience 7: 63. doi:10.3389/fnint.2013.00063

Elul R. (1969): Gaussian behavior of the electroencephalogram: changes during performance of mental task. Science 164: 328–331.

Fingelkurts A.A. & Fingelkurts A.A. (2010): Short-term EEG spectral pattern as a single event in EEG phenomenology. The Open Neuroimaging Journal 4: 130–156.

Fingelkurts A.A. & Fingelkurts A.A. (2015): Operational architectonics methodology for EEG analysis: theory and results. Neuromethods 91: 1–59. doi:10.1007/7657_2013_60

Freeman W.J. (1964): A linear distributed feedback model for prepyriform cortex. Exptl. Neurol. 10: 525–547.

Freeman W.J. (1967): Analysis of function of cerebral cortex using systems theory. Logistics Rev. 3: 5–40.

Freeman W.J. (1968): Relations between unit activity and evoked potentials in prepyriform cortex of cats. Journal of Neurophysiology 31: 337–348.

Garrido M.I., Kilner J.M., Kiebel S.J. & Friston K.J. (2007): Evoked brain responses are generated by feedback loops. PNAS 104: 20961–20966.

Güntekin B. & Başar E. (2016): Review of evoked and event-related delta responses in the human brain. International Journal of Psychophysiology 103: 43–52.

Hari R. & Parkkonen L (2015): The brain timewise: how timing shapes and supports brain function. Phil. Trans. R. Soc. B 370: 20140170

Hashemi P., Dankoski E.C., Lama R., Wood K.M., Takmakov P. & Wightman R.M. (2012): Brain dopamine and serotonin differ in regulation and its consequences. PNAS 109: 11510–11515.

Hoeks B. & Ellenbroek B.A. (1993): A neural basis for a quantitative pupillary model. Journal of Psychophysiology 7: 315–324.

Hutchison R.M., Womelsdorf T., Allen E.A. et al. (2013): Dynamic functional connectivity: promise, issues, and interpretations. Neuroimage 80: 360–378. doi: 10.1016/j.neuroimage.2013.05.079

Inouye T., Toi S. & Matsumoto Y. (1995): A new segmentation method of electroencephalograms by use of Akaike’s information criterion. Brain Res. Cogn. Brain Res. 3: 33–40.

Intriligator J. & Polich J. (1995): On the relation between EEG and ERP variability. International Journal of Psychophysiology 20: 59–74.

Jia H., Hu X. & Deshpande G. (2014): Behavioral relevance of the dynamics of the functional brain connectome. Brain Connectivity 4: 741–759. doi: 10.1089/brain.2014.0300

Kaiser R.H., Whitfield-Gabrieli S., Dillon D.G., Goer F., Beltzer M., Minkel J. et al. (2015): Dynamic resting-state functional connectivity in major depression. Neuropsychopharmacology 41: 1822–1830. doi: 10.1038/npp.2015.352

Kaplan A. Ya. (1999): The problem of segmental description of human electroencephalogram. Human Physiology 25: 107–114.

Klados M.A., Frantzidis C., Vivas A.B., Papadelis C., Lithari C., Pappas C. & Bamidis P.D. (2009): A framework combining delta event-related oscillations (EROs) and synchronization effects (ERD/ERS) to study emotional processing. Computational Intelligence and Neuroscience 2009 (Article ID 549419).

Knyazev G. G. (2007): Motivation, emotion, and their inhibitory control mirrored in brain oscillations. Neuroscience and Biobehavioral Reviews 31: 377–395.

Knyazev G.G. (2012): EEG delta oscillations as a correlate of basic homeostatic and motivational process. Neuroscience and Biobehavioral Reviews 36: 677–695.

Labuschagne I., Croft R.J., Phan K.L. & Nathan P.J. (2010): Augmenting serotonin neurotransmission with citalopram modulates emotional expression decoding but not structural encoding of moderate intensity sad facial emotional stimuli: an event-related potential (ERP) investigation. Journal of Psychopharmacology 24: 1153–1164.

Lavin C., San Martin R. & Rosales Jubal E. (2014): Pupil dilation signals uncertainty and surprise in a learning gambling task. Frontiers in Behavioral Neuroscience 7: 218. doi: 10.3389/fnbeh.2013.00218

Lieder F., Stephan K.E., Daunizeau J., Garrido M.I. & Friston K.J. (2013): A neurocomputational model of the mismatch negativity. PLOS Computational Biology 9: e1003288.

Makeig S. & Onton J. (2009): ERP Features and EEG Dynamycs: An ICA Perspective. In: S. Luck & E. Kappenman (Eds.): Oxford Handbook of Event-Related Potential Components. New York, Oxford University Press.

Muller A., Joseph V., Slesinger P.A. & Kleinfeld D. (2014): Cell-based reporters reveal *in vivo* dynamics of dopamine and norepinephrine release in murine cortex. Nature Methods 11: 1245–1252.

Nieuwenhuis S., Aston-Jones G. & Cohen J.D. (2005): Decision making, the P3, and the Locus Coeruleus-Norepinephrine System. Psychological Bulletin 131: 510–532.

Nieuwenhuis S., De Geus E.J. & Aston-Jones G. (2011): The anatomical and functional relationship between the P3 and autonomic components of the orienting response. Psychophysiology 48: 162–175.

Olofsson J.K., Nordin S., Sequeira H. & Polich J. (2008): Affective picture processing: an integrative review of ERP findings. Biological Psychology 77: 247–265.

Petridou N., Gaudes C.C., Dryden I.L., Francis S.T. & Gowland P.A. (2012): Periods of rest in fMRI contain individual spontaneous events which are related to slowly fluctuating spontaneous activity. Human Brain Mapping 34:1319-1329. doi: 10.1002/hbm.21513

Polich J. (2007): Updating P300: An integrative theory of P3a and P3b. Clinical Neurophysiology 118:2128-2148.

Preuschoff K., ’t Hart B.M. & Einhäuser W. (2011): Pupil dilation signals surprise: evidence for noradrenaline’s role in decision making. Frontiers in Neuroscience 5: 115. doi: 10.3389/fnins.2011.00115

Rashid B., Damaraju E., Pearlson G.D. & Calhoun V.D. (2014): Dynamic connectivity states estimated from resting fMRI identify differences among schizophrenia, bipolar disorder, and healthy control subjects. Front. Hum. Neurosci. 8: 897. doi: 10.3389/fnhum.2014.00897

Roschke J. & Fell J. (1997): Spectral analysis of P300, generation in depression and schizophrenia. Neuropsychobiology 35: 108–114.

Schürmann M., Başar-Eroglu C., Kolev V. & Başar E. (2001): Delta responses and cognitive processing: single-trial evaluations of human visual P300. International Journal of Psychophysiology 39: 229–239.

Staib M., Castegnetti G., Bach DR (2015): Optimising a model-based approach to inferring fear learning from skin conductance responses. Journal of Neuroscience Methods 255: 131–138.

Stampfer H.G. & Başar E. (1985): Does frequency analysis lead to better understanding of human event-related potentials? Int. J. Neurosci. 26: 181–196.

Tagliazucchi E., Balenzuela P., Fraiman D. & Chialvo D.R. (2012): Criticality in large-scale brain fMRI dynamics unveiled by a novel point process analysis. Frontiers in Physiology 3:15. doi: 10.3389/fphys.2012.00015

Teplan M. (2002): Fundamentals of EEG measurement. Meas Sci Rev 2: 1–11.

Vaughan H. G. Jr. (1969): The relationship of brain activity to scalp recordings of event-related potentials. In: E. Donchin & D.B. Lindsly (Eds.), Averaged evoked potentials: Methods, results, evaluations. Washington, D.C.: NASA, pp. 45–94.

Venkatraman A., Edlow B.L. & Immordino-Yang M.H. (2017): The brainstem in emotion: a review. Front. Neuroanat. 11: 15. doi: 10.3389/fnana.2017.00015

Woodman G.F (2010): A brief introduction to the use of event-related potentials in studies of perception and attention. Attention, Perception & Psychophysics 72: 2031–2046.

